# Development of metagenomic methods for non-invasive health monitoring of endangered species: Unveiling hidden microbial threats in fecal samples

**DOI:** 10.1101/2025.01.21.633432

**Authors:** Román Sapino, Ángel Fernández-González, Jose Castresana

## Abstract

Metagenomic analysis of feces is emerging as a powerful tool for improving the monitoring of endangered species. A critical aspect in assessing the extinction risk of a species is the analysis of the burden of parasites and pathogens that can negatively affect the health of individuals. However, the identification of pathogens in non-model species using metagenomics is a major challenge due to the lack of reference genome sequences or data limited to distantly related species. In this study, we developed a pipeline for detecting potentially pathogenic bacteria from metagenomic sequences by mapping unassembled reads to available reference genomes. The approach uses the breadth of genome coverage rather than the number of mapped reads for species identification, thereby minimizing false positives due to conserved or repetitive genomic regions. We applied this method to fresh fecal samples of the Iberian desman (*Galemys pyrenaicus*), a critically endangered semi-aquatic mammal. Our analysis revealed the presence of 19 potentially pathogenic bacterial species, with prevalences ranging from a single individual to 30% of the samples. We also detected some desmans with elevated or altered pathogen loads, suggesting variations in individual health status or different environmental exposures. This work represents a novel application of metagenomic methods for species-level pathogen detection in wildlife using fecal samples. Application of this method across populations and over time for endangered species may provide essential health and epidemiological information to improve conservation strategies.

## Introduction

Parasitic and pathogenic species can have a significant impact on the survival and reproduction of wild species (Daszak et al., 2000; De Castro & Bolker, 2004; Smith et al., 2006; Pedersen et al., 2007; Blanchong et al., 2016). In the case of endangered species, understanding the burden and diversity of these pathogens is essential for developing effective conservation strategies. Traditional methods of studying infectious diseases often require invasive sampling techniques, which can be particularly problematic for endangered species. In contrast, non-invasive methods, such as fecal analysis or environmental DNA, offer a valuable alternative to gather vital health information without disturbing the animals (Beja-Pereira et al., 2009; Queirós et al., 2023).

Metagenomics has revolutionized our ability to profile the taxonomic composition of complex samples, including those present in fecal samples. This advance has made it possible to study areas as diverse as diet, the microbiome, and even the host genome using only feces (Srivathsan et al., 2015; Srivathsan et al., 2016; Quince et al., 2017; Gibson et al., 2019; Taylor et al., 2022; de Flamingh et al., 2023). Nevertheless, despite the potential of metagenomics to provide detailed insights into the presence and diversity of parasites and pathogens using fecal samples, its application to the accurate detection of these species and the health assessment of endangered species remains largely unexplored since it was first proposed (Srivathsan et al., 2015; Srivathsan et al., 2016).

Several metagenomic methods area available to detect pathogenic bacteria from environmental samples such as feces. The programs Kraken (Lu et al., 2022) and Diamond (Buchfink et al., 2015) employ lowest common ancestor strategies to provide taxonomic classification for as many reads as possible. These methods accurately identify species when reference sequences are available, or otherwise provide higher-order classification. They are particularly popular in microbiome studies where the focus is often on community composition at higher taxonomic levels (Quince et al., 2017; Gibson et al., 2019; Pinto & Bhatt, 2024). Assembly-based methods, on the other hand, require high coverage to assemble genomes and may only detect a fraction of the species in a sample due to the depth of sequencing required, being less effective for low abundance species and complex microbial communities (Nurk et al., 2017; Blanco-Miguez et al., 2023). Furthermore, several pipelines have been specifically developed for detecting pathogenic species in clinical applications. These include PathoScope (Francis et al., 2013; Hong et al., 2014) and SURPI (Naccache et al., 2014; Gu et al., 2021). These pipelines use as main engine mapping programs designed for resequencing studies, where reads are aligned to a reference genome. PathoScope utilizes Bowtie2 (Langmead & Salzberg, 2012), while SURPI employs SNAP (Zaharia et al., 2011) for mapping. Kraken also recommends mapping with Bowtie2 for corroborating pathogen identification at the species level (Lu et al., 2022). Mapping methods are optimized for speed in whole genome analysis, which can only be achieved by mapping reads to the reference genomes of the same or very closely related species. These methods also calculate a mapping quality (MAPQ) score (Li et al., 2008; Langmead, 2017), which reflects the probability that a read is aligned to the correct position in the reference genome. Consequently, MAPQ also serves as a proxy for the confidence that the read truly originates from the species genome used for mapping. Due to these characteristics, and although mapping methods were not initially developed for species identification, they have proven to be very useful for this purpose due to their efficiency and accuracy in aligning reads to specific reference genomes. Despite significant progress in methods for the accurate identification of pathogens and parasites in human health applications, there remains a notable gap in the application of these techniques to non-model species, mainly due to the need for reference complete genomes of the exact or closely related species in databases. However, the increasing number of complete genomes now supports their broader use (Goldfarb et al., 2024). Developing methods to obtain accurate genomic information about pathogens from feces is imperative for enhancing pathogen surveillance in endangered species.

The Iberian desman (*Galemys pyrenaicus*) is a critically endangered semi-aquatic mammal endemic to the Iberian Peninsula (Palmeirim & Hoffmann, 1983). The species faces numerous threats, including habitat destruction, water pollution, and the presence of barriers such as reservoirs and hydroelectric power plants that disrupt its riverine habitat (Quaglietta et al., 2024). These barriers not only fragment the desman’s habitat, but also isolate its populations, leading to significant inbreeding problems that further jeopardize the species’ survival (Escoda et al., 2019; Escoda et al., 2022). Despite its endangered status, there is a lack of comprehensive research on pathogens affecting the Iberian desman, with only studies using PCR-based methods to detect pathogens in this species (Ripa et al., 2023). However, populations of this species have been declining rapidly or disappearing over the last two decades for reasons that remain largely unknown. Understanding the impact of pathogens and parasites is essential not only for the management and recovery of Iberian desman populations *in situ*, but also for planning any future conservation strategies, such as captive breeding and translocations, as these strategies carry the risk of inadvertently spreading diseases if the health status of the populations involved is not thoroughly understood (Gaywood et al., 2022). Therefore, detailed studies of the pathogens and parasites affecting the Iberian desman are critical to ensure the success of conservation efforts, prevent further population extinctions, and aid in their recovery.

The aim of this study is to develop and test a pipeline based on mapping of unassembled reads to reference genomes to identify pathogenic bacterial species in feces using metagenomic sequencing. Fresh samples from the Iberian desman were used to test the method. Our approach specifically addresses several limitations of previous methods, particularly when applied to non-model species that lack comprehensive reference genomes of pathogens, and adapts them to these unique challenges. By focusing on bacteria, which have smaller genomes and are better represented in genomic databases, we aim to establish an efficient and reliable approach that can be used for long-term health monitoring of microbial pathogens in endangered species.

## Materials and Methods

### Fecal samples collection and DNA extraction

Fecal samples were collected in 2018 and 2019 from the Iberian desman population of the Central System, located in the center of the Iberian Peninsula. A total of 23 samples (Table S1) were obtained from four different hydrological units or subpopulations: Becedillas, Aravalle, Endrinal and Adaja. Fecal samples were collected during the capture of individuals immediately after deposition, and placed in tubes with ethanol. This approach preserves DNA integrity and minimizes the risk of environmental contamination that can occur with fecal samples (Hawlitschek et al., 2018; Oliveros et al., 2023). The work of capturing individuals was part of a conservation program independent of this study promoted by the Ministry of Environment through the Biodiversity Foundation, the Duero River Basin Authority, and the Autonomous Government of Castilla y León through the Patrimonio Natural Foundation, in Spain.

DNA was extracted using the QIAamp DNA Mini Kit (QIAGEN) following the manufacturer’s instructions and quantified using a Qubit fluorometer with the Qubit dsDNA High Sensitivity Assay Kit (Thermo Fisher Scientific).

### Shotgun metagenomic library construction and sequencing

Shotgun metagenomic libraries were constructed using the NEBNext Ultra II FS DNA Library Prep Kit (New England Biolabs). Extracted DNA (26 μL per sample) was enzymatically fragmented at 37°C for 15 minutes. Specific Illumina adapters were ligated, and fragments of 150-250 bp insert size were selected using NEBNext Sample Purification Beads (New England Biolabs). Each sample was then indexed and amplified through 12 cycles of PCR, cleaned, and quantified using the Qubit fluorometer. Fragment size distribution (270 – 370 bp) was assessed by E-gel EX 2% agarose gel electrophoresis (Invitrogen). Finally, equimolar amounts of each library were pooled and sequenced on an Illumina platform at Macrogen Inc. (South Korea).

### Quality control and filtering of endogenous sequences

Reads of low quality, shorter than 100 bp, or duplicated were filtered with FASTP version 0.23.2 (Chen, 2023), while sequences with repetitive motifs were filtered with BBDUK 39.01 (https://jgi.doe.gov/data-and-tools/software-tools/bbtools/). The remaining reads were aligned to the reference *G. pyrenaicus* genome (Escoda & Castresana, 2021) using Bowtie2 version 2.5.0 (Langmead & Salzberg, 2012). Unmapped sequences were used in subsequent steps to detect pathogenic bacteria.

### Reference bacterial genomes

Reference genomes for the bacterial species were obtained from the NCBI Genome Database (https://www.ncbi.nlm.nih.gov/datasets/). The genus *Yersinia* was detected in initial analyses of desman samples and was chosen as a case study to evaluate different methods for species identification. We specifically selected for testing a group of closely related species, which can present significant identification problems. The genus contains 26 species listed in the NCBI taxonomy, four of which are pathogenic (Table S2), so accurate identification to species level is essential in this genus, highlighting the need for robust methods for this task.

For a broader analysis of other pathogenic bacteria, the reference genomes of species found in the Pathogen Detection Project of NCBI (https://www.ncbi.nlm.nih.gov/pathogens/) and the Virulence Factor Database (VFDB, http://www.mgc.ac.cn/VFs/) (Liu et al., 2022) were downloaded, in addition to other bacterial species found in the literature (Barandika et al., 2007; Pedersen et al., 2007; Cantas & Suer, 2014; White & Razgour, 2020; Ali & Alsayeqh, 2022; Sabour et al., 2022; Suminda et al., 2022) and online sources (e.g. https://ewda.org/diagnosis-cards/, animal health section of https://www.mapa.gob.es, etc.), resulting in 137 species that included the four pathogenic *Yersinia* species (Table S3). *Cutibacterium acnes* was included in the initial analyses and was found in most samples, but it has been reported that this species is a likely contaminant of kits and reagents (Gu et al., 2019) and was excluded from the final analyses, which were based on 136 species.

### Alignment of sequences to reference genomes of *Yersinia* species

The set of 26 *Yersinia* species was used to test the performance of different methods and parameters in the pipeline. First, the set of exogenous reads was aligned separately to each of the respective reference genomes using Bowtie2 (Langmead & Salzberg, 2012) in the end-to-end alignment mode and the “--sensitive” option. In addition, the options “--no-discordant” and “--no-mixed” were used to capture only read pairs where both reads align and have the expected orientation and distance between them. Subsequently, alignments were converted to BAM format, PCR duplicates removed, and reads with MAPQ values lower than 20 filtered using SAMtools v1.9 (Li et al., 2009). General mapping quality was visualized using the coverage across the genome graph constructed with Qualimap 2.2.2 (Okonechnikov et al., 2016), which represents the mean depth of coverage in 4,000 windows of the genome.

Using SAMtools, genome alignment statistics were calculated for each sample and bacterial species, including the number of mapped reads, the mean MAPQ of the mapped reads, the number of positions in the reference genome sequenced with at least one read, the breadth of coverage (%; percentage of the reference genome covered by at least one read), and the depth of coverage (X; mean number of mapped reads at each genome position). Different thresholds based on percent coverage were set to determine the presence of each species in the samples. We also calculated the percent coverage when filtering out positions with extreme values of depth of coverage, defined as those sites with a depth of coverage greater than the sample mean plus 3 times the standard deviation, calculated in logarithmic space.

Bowtie2 can be used with single-species or multi-species databases. A Bowtie2 database was first constructed for each species for the main analyses. In addition, a combined database containing the set of all species was constructed to assess the effect of database configuration on analysis performance. For samples with multiple positives of species in the same genus or closely related species, the numbers of unique and shared reads for each positive species were compared using Venn diagrams.

The set of exogenous sequences was also aligned against the *Yersinia* reference genomes using other methods and conditions. Thus, we used Bowtie2 in local alignment mode, as well as another widely used DNA mapping program, BWA 0.7.17 (Li & Durbin, 2009). For the latter, both the BWA-mem and BWA-aln algorithms were evaluated in their default configurations. The MAPQ scales are different in Bowtie2 and BWA, with a maximum of 42 for Bowtie2 and 60 for BWA. As mentioned above, a MAPQ cut-off of 20 was used for Bowtie2. The distributions of MAPQ values for both methods in the same set of samples showed that the equivalent threshold for BWA was ∼30, so this value was used to filter mapped reads with this program.

### Alignments to reference genomes of pathogenic bacterial species

The Bowtie2 end-to-end mode, which provided consistent results for the *Yersinia* species, was used to align the exogenous reads from each sample to the 136 pathogenic bacterial species using both single-species and combined databases. Identification was based on the percent coverage thresholds as described above. Principal Component Analyses (PCA) was performed on the percent coverage values to compare the overall health status of the Iberian desmans using the single-species databases (similar results were obtained using the combined database).

## Results

### Species-level identification of *Yersinia*

From the 23 fecal samples, we obtained between ∼30 and ∼80 million exogenous reads per sample after eliminating Iberian desman sequences (Table S1). To evaluate the mapping approach for identifying closely related bacterial species, we first used the Bowtie2 aligner (end-to-end mode) to map reads from each sample to the genomes of 26 *Yersinia* species (Table S4), with a Bowtie2 database constructed per genome.

Traditional species identification often relies on the number of mapped reads. However, coverage plots across the genome revealed that certain regions, either because they are highly conserved among different species in the sample or because they are repetitive, tend to accumulate a large number of reads, while the rest of the genome maps any read. This is illustrated in Figure 1, which compares the mappings of two samples, BC3903 and BC3345, to the *Yersinia aldovae* genome. Sample BC3903, with 12,378 reads, showed a mostly even distribution of reads across the genome, as expected for the mapping of reads in the genome of the same species. In contrast, BC3345, with a similar number of reads (13,790), showed extremely high coverage in certain regions (up to ∼250X), while most regions of the genome were not covered by reads, suggesting that *Y. aldovae* is not present in this sample. Similar contrasting examples are shown in Figure S1. Therefore, direct application of the number of mapped reads to species identification may lead to false positives.

**Figure 1.**
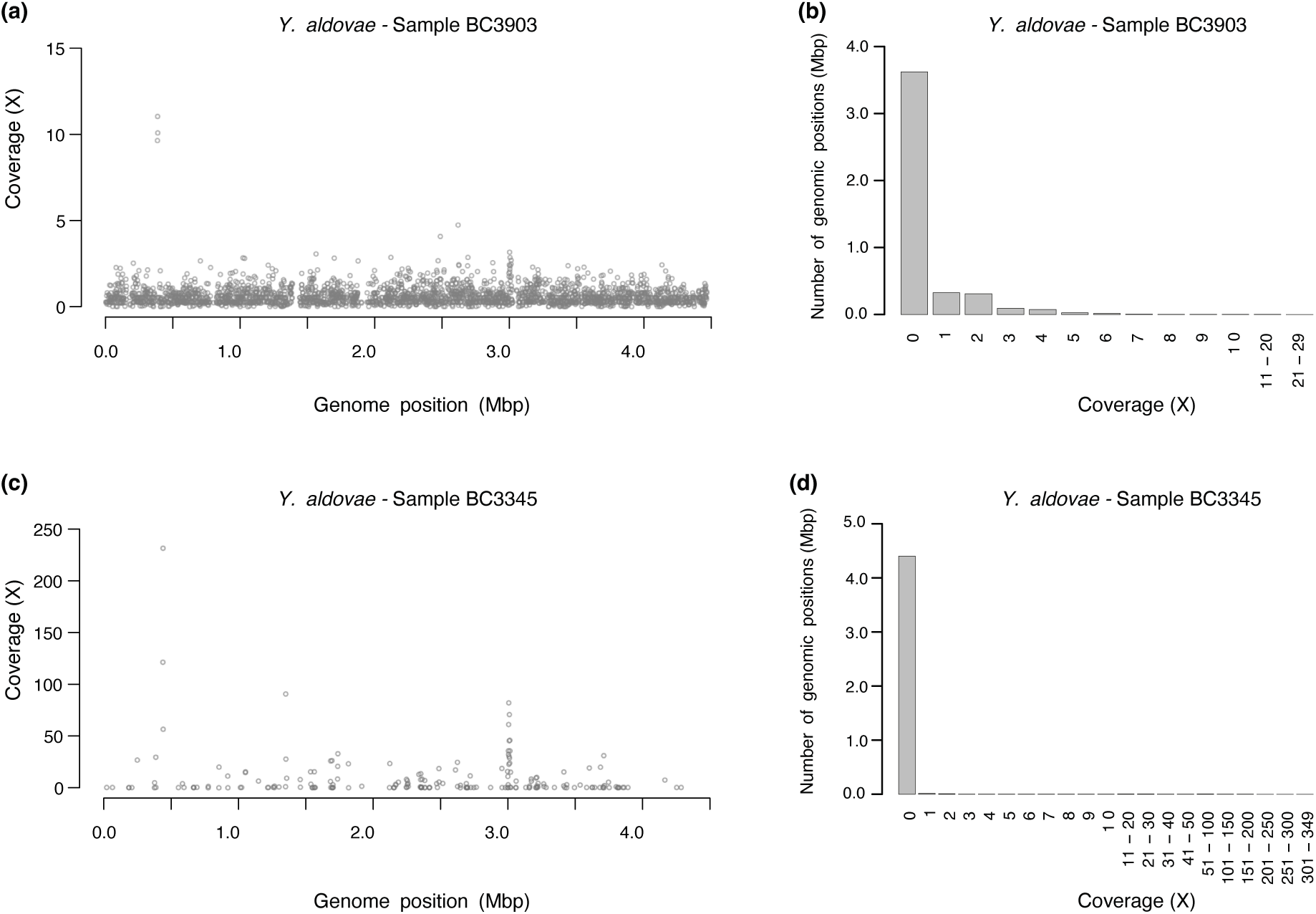
Comparison of depth of coverage across the reference genome of *Yersinia aldovae* and the corresponding depth of coverage barplots for two samples of Iberian desman. (a) Coverage across the reference genome in 4,000 windows and (b) coverage barplot for the BC3903 sample, in which 12,378 reads were assigned, resulting in a mean depth coverage of 0.41X and a breadth of coverage of 19.03%. (c) Coverage across the reference genome and (d) coverage barplot for the BC3345 sample, in which 13,790 reads were assigned, resulting in a mean depth coverage of 0.46X and a breadth of coverage of 1.5%. In (a) and (c), windows with zero coverage are not plotted. Note the different coverage scales in (a) and (c).

Given the large number of unspecific reads mapped, we found that the breadth of coverage was a more reliable parameter for species identification than the total number of reads. For example, in the previous example, sample BC3903 had 19.03% of its genome covered by at least one read, strongly suggesting the presence of *Y. aldovae* or a close relative, whereas BC3345 had only 1.5% covered (Figure 1 and Table S4), despite having a similar number of mapped reads. However, setting a threshold for the percent coverage for positive identification is not straightforward. According to initial analyses of *Yersinia* and other species, we explored the performance of two different thresholds for species identification: a first threshold at ≥3% coverage and a second at ≥2% coverage (corresponding to ∼139 kb and 92 kb of an average *Yersinia* genome covered, respectively).

Using the stricter 3% threshold (Figure 2a), we identified 13 instances of 5 distinct *Yersinia* species. *Y. intermedia* had the highest prevalence, being detected in 9 samples, while *Y. aldovae*, *Y. entomophaga*, *Y. nurmii*, and *Y. ruckeri* were found in only one sample each. Percent coverages of these 13 identifications ranged from 3.35 to 45.38% of the genome, which perfectly correlated with number of mapped reads (Table S4). The mean MAPQ values for the mapped reads of the positive identifications were close to the maximum value (ranging from 40.66 to 41.89), meaning that most reads were perfectly aligned in the correct position in the reference genome used. Lowering the threshold to 2% genome coverage resulted in three additional identifications (Figure 2a). The previously found species *Y. intermedia* was detected in one additional sample, and two new species, *Y. alsatica* and *Y. enterocolitica*, were detected in only one sample each. Similar identification results were obtained when using a combined Bowtie2 database with all *Yersinia* species (Figure S2a). However, three of the detections were missing, likely indicating fewer unspecific mapped reads with combined databases. We also verified that the percent coverage was very similar when positions with extreme depth of coverage were filtered out, resulting in the same identifications.

**Figure 2.**
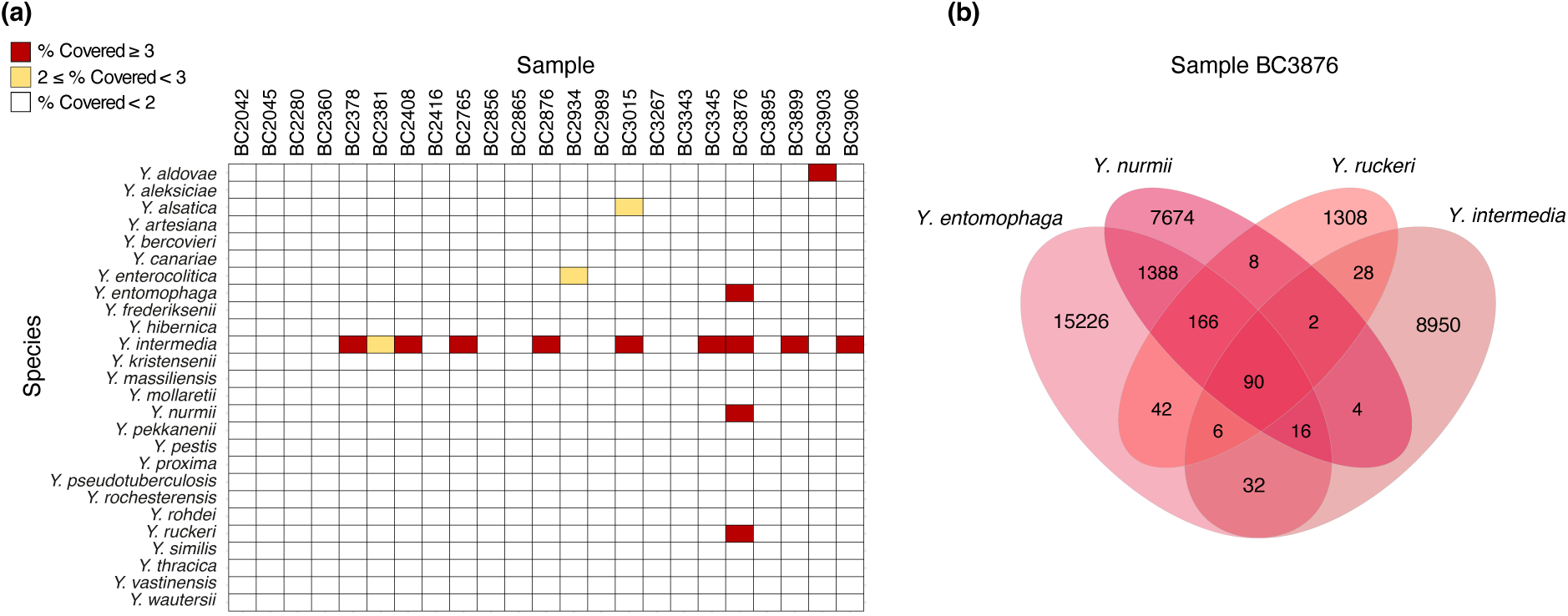
Identification of *Yersinia* species in 23 samples of Iberian desman and evaluation of read sharing of multiple species found in a single sample. (a) Species identification according to the percentage of the genome covered by mapped reads. (b) Venn diagram comparing reads assigned to four *Yersinia* species in sample BC3876, showing shared reads (intersections) and unique reads (non-intersecting areas) for each species.

In sample BC3876, where four *Yersinia* species were identified, we used a Venn diagram to compare mapped reads and assess whether all four species were actually present in the sample. We found that each species had a significant number of unique reads, with the only exception of *Y. entomophaga* and *Y. nurmii*, which shared a substantial number of reads (1,660, representing 18% of the reads mapped to *Y. nurmii*; Figure 2b). The analysis with the combined database including all *Yersinia* species corroborated this finding, as *Y. nurmii* was not positive in BC3876 (Figure S2a). However, all other three species were consistently detected in the same sample in both analyses, which, together with the low number of shared reads among the three species, suggests that all three are likely present in the sample.

### Comparison of mapping tools for species identification

To verify the mappings obtained with the default Bowtie2 end-to-end algorithm, we compared them with those generated using Bowtie2 in local mode, BWA-mem, and BWA-aln. For all samples, BWA-mem reached high mapping rates (e.g., an average of 270,298 reads aligned to *Y. intermedia*, compared to 6,511, 6,459, and 7,540 reads for Bowtie2 end-to-end, Bowtie2 local mode, and BWA-aln, respectively) and also higher breadth of coverage values. This was achieved mostly by reducing both read length and insert size, indicating that this mapping algorithm, in its default configuration, exhibits less stringent behavior and has no discrimination power at the species level. As for the other three algorithms, the number of reads mapped to each sample and the breadth of coverage values were similar (Figure 3 shows the results for *Y. intermedia*). BWA-aln achieved a higher number of mapped reads, although a consistent pattern was maintained for the three methods across all samples. Consequently, the determination of the *Yersinia* species was similar for these three methods, with both Bowtie2 algorithms giving identical identifications and BWA-aln giving some additional positives (Figures S2b and S2c). However, the results would also be equivalent for the latter method if the breath of coverage thresholds used for positive identification were raised slightly. Therefore, the three stringent mapping algorithms showed similar performance in species identification.

**Figure 3.**
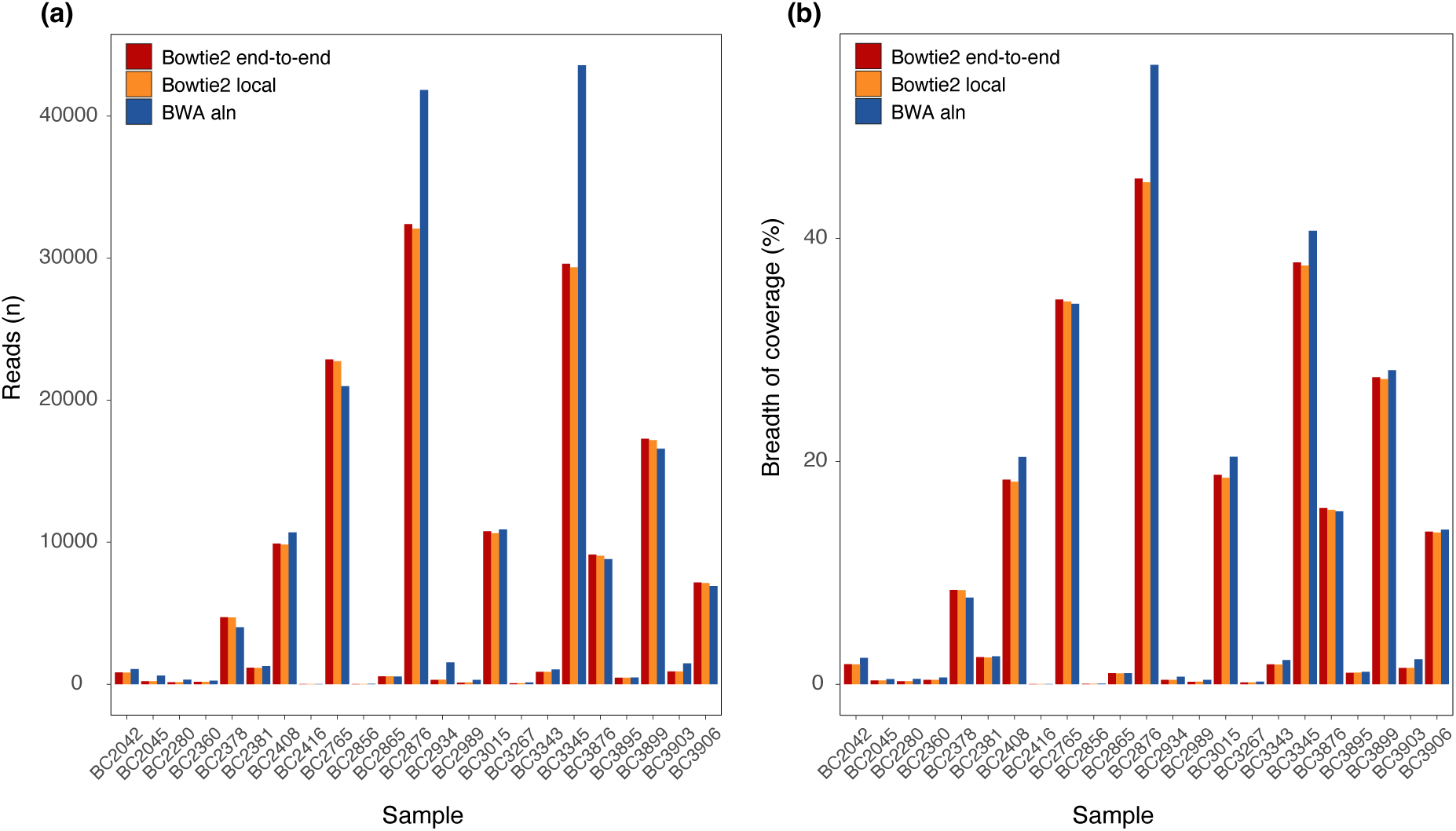
Comparison of different mapping tools (Bowtie2 end-to-end, Bowtie2 local, and BWA-aln) using the *Yersinia intermedia* genome as an example. (a) Number of reads aligned to each sample by each mapping tool. (b) Percent of the genome covered achieved by each mapping tool.

### Application to the detection of pathogenic bacterial species

Due to its consistent performance, the Bowtie2 end-to-end algorithm was used in conjunction with the single-species databases for the pathogenic bacteria identification. Using the more stringent 3% genome coverage threshold, we identified 57 instances of pathogenic species belonging to 19 different species (Figure 4). Table 1 details these 19 bacterial species, their associated pathologies, and other relevant wildlife species in which these pathogens were found. The most frequently detected species were *Acinetobacter baumannii* and *Fusobacterium necrophorum*, each found in 7 samples. Samples BC2381 and BC2934 had the highest number of pathogenic species, with 14 and 8 species, respectively. Notably, all but 7 samples were found to contain at least one pathogen. Percent coverages started at 3.29% and were as high as 91.74% for *Shigella sonnei* in one sample. The mean MAPQ values for the positive identifications were close to the maximum (ranging from 40.95 to 41.98). When the threshold was lowered to 2% coverage, 19 additional identifications were obtained. However, many of these corresponded to species detected in only one sample (Figure 4). A similar identification pattern was observed when all genomes were analyzed using a combined Bowtie2 database, although the total number of identifications was reduced to 50 with the 3% threshold, and only 7 additional identifications were obtained with the 2% threshold (Figure S3).

**Figure 4.**
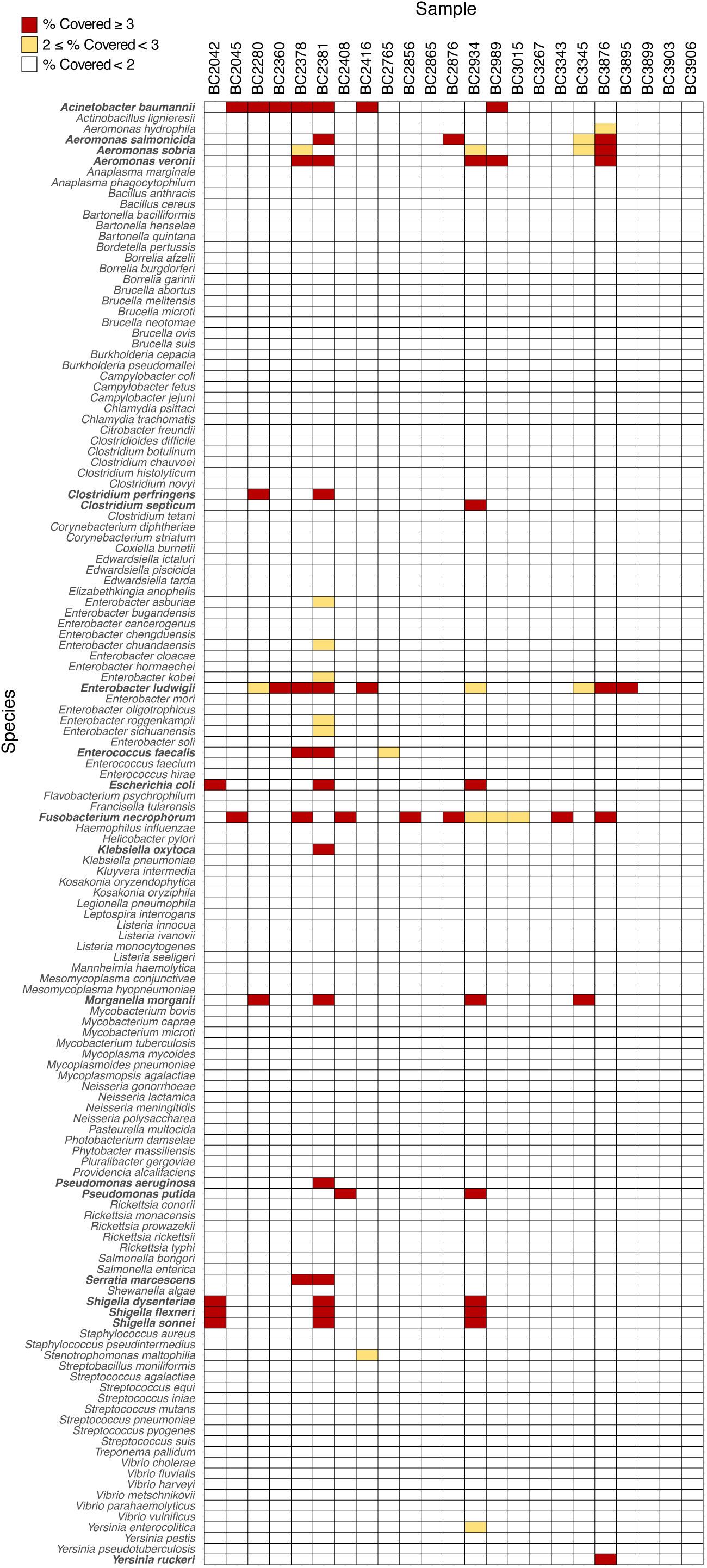
Identification of 136 bacterial species in 23 samples of Iberian desman. Species were identified according to the percentage of the genome covered by mapped reads. Species with coverage greater than 3% are shown in bold for clarity.

**Table 1.**
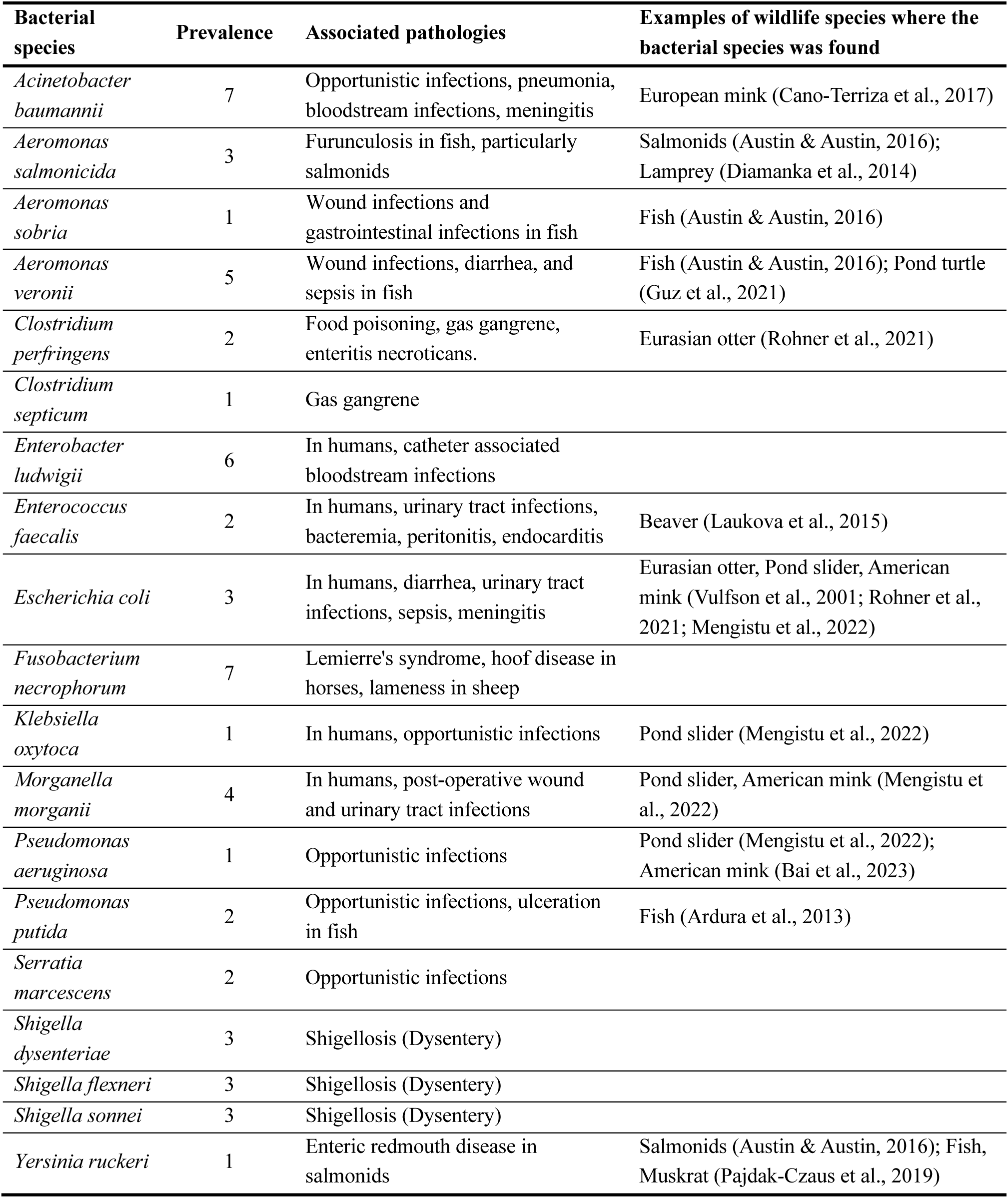
Bacterial species found in Iberian desman feces with the threshold of 3% of the genome covered by mapped reads. For each species, its prevalence and a brief description of associated pathologies as found in different sources are given. In addition, examples of other relevant wildlife species in which the bacterial species has been found are given.

In four cases where species of the same genus or closely related species were detected in a single sample, we used Venn diagrams to check species identification accuracy. For instance, three *Aeromonas* species were detected in sample BC3876 with each having a significant number of unique reads, confirming the likely presence of the three species in this sample (Figure S4). Samples where only one or two of the three species were assigned support this result. Conversely, *Escherichia coli* and the three species of the genus *Shigella*, which are four very closely related species (Chattaway et al., 2017), coincided in three samples, but the Venn diagram shows that these species share most reads (Figure S4). When all genomes were analyzed in a combined database, only one sample was positive for these four species whereas the other two samples were negative for the four bacteria (Figure S3). Thus, in analyses of very closely related species, read sharing due to unspecific mappings is found when using separate databases, whereas loss of read mappings due to ambiguous mapping and low MAPQ values occur in combined databases.

### Overall health assessment of the Iberian desman population

A PCA based on the breadth of coverage showed that most of the Iberian desman samples clustered together, with several outliers clearly defined (Figure 5). In particular, samples BC2381 and BC2934, previously mentioned as having the highest number of pathogenic bacterial species, are appreciably separated from the main group. Other outliers in the plot (BC3876, BC2045 and BC2378) show varying numbers of assigned pathogenic species, although they generally had a high number of pathogens with a high breadth of coverage. Four of the desmans with atypical values were from the Endrinal, while one belonged to the Adaja hydrological unit (Table S1).

**Figure 5.**
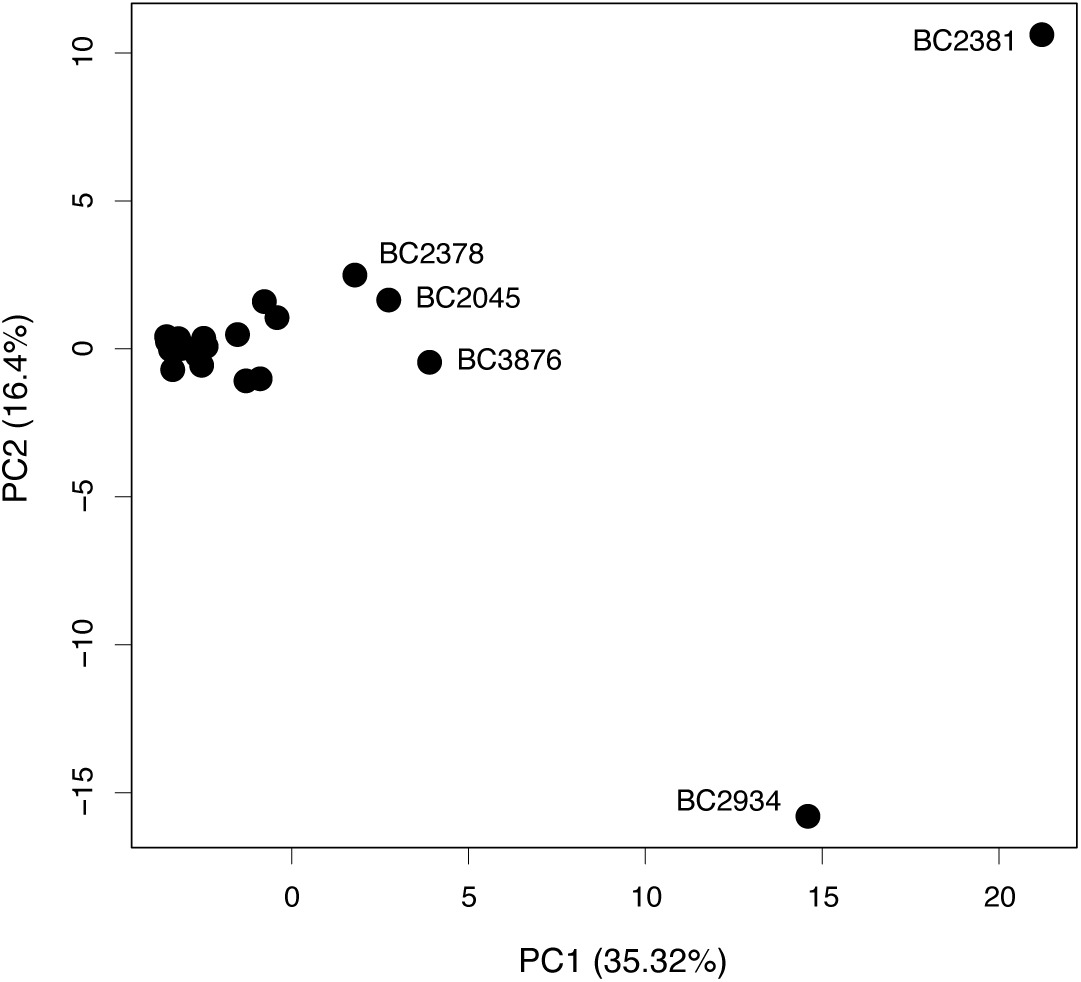
Principal Component Analysis based on breadth of coverage of each pathogenic bacterial species identified in 23 Iberian desman samples. The sample names of outliers are indicated. The percentage of variance explained by each principal component (PC1 and PC2) is shown on the corresponding axis.

## Discussion

### Validation of a mapping methodology for the detection of pathogens from fecal samples

In this work, we developed and validated a method to identify bacterial pathogens from fecal samples based on the mapping of unassembled reads obtained by metagenomic sequencing to reference genomes. This approach differs fundamentally from methods designed to profile the general composition of the microbiome, such as Kraken (Lu et al., 2022) or Diamond (Buchfink et al., 2015). Instead, it aligns more closely with pathogen detection methods used in human disease studies, where the primary objective is species-level identification of target pathogens, such as PathoScope (Francis et al., 2013; Hong et al., 2014) or SURPI (Naccache et al., 2014; Gu et al., 2021). To our knowledge, no studies have systematically applied these species-level pathogen detection methods to endangered wildlife species.

The effectiveness of our pipeline for wildlife health monitoring was demonstrated by its ability to identify pathogenic bacterial species in fecal samples from the Iberian desman. Several factors affecting the identifications were analyzed to better understand their impact, optimize the method, and guide their application in different scenarios. These factors included the choice of identification parameters, threshold settings, database configuration, and mapping tool.

To improve the accuracy of species identification, we used the breadth of genome coverage rather than the more commonly used number or proportion of mapped reads (Francis et al., 2013; Hong et al., 2014; Naccache et al., 2014; Gu et al., 2021), thus avoiding false positives caused by conserved or repetitive genomic regions, where reads from different species may map. Unspecific mappings can be particularly significant when the genome of the exact bacterial species present in the sample is not in the database and rather a closely related species is used for mapping. Therefore, using breadth of coverage for identification is especially important in studies of non-model and wildlife species, where the representation of pathogens in databases is often incomplete.

We analyzed two different operational thresholds for identifications using breath of coverage, corresponding to 2% and 3% coverage, to evaluate their sensitivity and specificity. In general, the more stringent 3% threshold identified species that were present in multiple samples, each with different genome coverage values that correlated with the number of mapped reads and, by extension, with species abundance (e.g., *Yersinia intermedia*, Table S4). This scaling of genome coverage with the number of reads strongly supports correct mapping to the reference genome and therefore that the species is truly present in the population. Conversely, species identified with breath of coverage values between 2 and 3% (or at lower levels) were typically detected in only one individual (Figures 2a and 4). Single-sample detections lacked the supporting evidence provided by the scaling of breath of coverage with the number of reads and, while their presence cannot be entirely ruled out, it is more difficult to dismiss for them the possibility of mappings to conserved or repetitive genomic regions of the genome of a distantly related species. Therefore, a 3% threshold provided more confident identifications in this particular study, given the sequencing effort applied. However, it is important to note that the most appropriate threshold may vary depending on the sequencing depth and specific conditions of individual studies. For instance, in cases involving highly pathogenic species or potential outbreaks, all samples, even those with low numbers of mapped reads, should be considered for further investigation to ensure comprehensive monitoring and risk assessment.

The type of database to be used with the mapping algorithm is another critical factor and its choice depends on the objectives of the study. For example, for targeted monitoring of highly pathogenic species, it would be essential to first perform an analysis of unspecific mappings (Figures 1 and S1) and shared read levels (Figures 2b and S4) for the pathogen of interest in relation to closely related species using different databases. Our results showed that unspecific mappings from sequences mapping to phylogenetically closely related species in the database, particularly in conserved regions, may be higher when using a single Bowtie2 database for each genome analyzed compared to the combined database. Furthermore, unspecific mappings may lead to more shared reads when using single-species databases. Despite this, single-species databases may be preferable for initial surveys, as using a combined Bowtie2 database for all genomes may result in the loss of potential positives due to ambiguous mappings with low mapping quality values, as observed for the *Shigella* and *Escherichia* species (Figures 4 and S3). In essence, single-species databases typically provide greater sensitivity because they focus on detecting specific pathogens without competition from sequences of other species, whereas combined databases improve specificity by offering a broader reference to discriminate between similar species, although they may reduce sensitivity due to ambiguous or low-confidence mappings to multiple species. Therefore, preliminary analyses should be conducted to determine the most suitable database and detection criteria for each pathogen under study to ensure that the method provides robust identifications.

The mapping tools Bowtie2 (in two different modes: local and end-to-end) and BWA (in aln mode) produced similar results in terms of number of mapped reads and breath of coverage achieved (Figure 3). This was not the case with the less stringent BWA program in mem mode, which uses a less stringent algorithm. The fact that Bowtie2 and BWA in their optimal configurations consistently produced similar mapping results and thus similar species identifications, despite their algorithmic differences, reinforces confidence in their ability to provide accurate taxonomic classifications.

The analysis of the genus *Yersinia* illustrates the efficiency of our approach. Of the 26 species tested, seven different species were detected in the Iberian desman samples when using the more comprehensive 2% threshold. Among these, two species (*Y. enterocolitica* and *Y. ruckeri*) are potentially pathogenic, with each found in a single sample (Figures 2 and 4). These results highlight the importance of robust species-level identifications and demonstrate the power of the method proposed here to discriminate between species within the same genus. However, challenges remain when resolving exact species in groups of very closely related taxa, such as *Shigella* and *Escherichia* species, where shared genomic regions can lead to ambiguous mappings. In these cases, assembly-based approaches combined with high-depth sequencing may be necessary to achieve accurate species-level resolution (Nurk et al., 2017; Blanco-Miguez et al., 2023).

### Implications for the conservation of the Iberian desman

Understanding pathogen burden and diversity in endangered species is critical to inform management strategies, particularly in the context of conservation efforts such as captive breeding and translocation programs (Gaywood et al., 2022). In our study, we identified 19 species of pathogenic bacteria in 23 Iberian desman fecal samples using a strict coverage threshold, with varying levels of genome coverage and prevalence. The sample size was very limited, which prevents any generalization of the results. Undoubtedly, a larger sample size, covering a wider geographical area and different time points, would provide a more comprehensive understanding of pathogen prevalence and diversity in this mammal. Although the main focus of the work was methodological, with the aim of developing a strategy that can be used for health monitoring, this pilot analysis also provided essential baseline data on the presence and diversity of pathogens in the Iberian desman, which could help shed light on the causes of the rapid decline of some of its populations. Remarkably, as shown in Table 1, many of the identified pathogenic species are found in aquatic environments and are known to infect fish (Ardura et al., 2013; Diamanka et al., 2014; Austin & Austin, 2016; Pajdak-Czaus et al., 2019) and other aquatic or semi-aquatic species, including European mink, American mink, Eurasian otter, beaver, muskrat, pond slider, and pond turtle (Vulfson et al., 2001; Laukova et al., 2015; Cano-Terriza et al., 2017; Pajdak-Czaus et al., 2019; Guz et al., 2021; Rohner et al., 2021; Mengistu et al., 2022; Bai et al., 2023). In some cases, these infections have been associated with disease or mortality in these animals, underscoring the potential severity of some of these pathogens. The sharing of pathogens among aquatic species may be explained by the fact that the aquatic environment can facilitate the transmission of infections, spreading pathogens over a wide area and potentially increasing exposure to multiple microbial threats (Cabral, 2010). Thus, these results highlight the complex and potentially pathogenic microbial landscape to which the Iberian desman is exposed.

The PCA (Figure 5) revealed substantial variability in pathogen profiles among Iberian desmans, suggesting differences in pathogenic bacterial load or diversity of species present, which may be highly relevant for assessing the health status of the analyzed individuals. In particular, five desmans emerged as clear outliers in the PCA plot, indicating altered pathogen loads. It remains unclear whether these desmans contained these levels of pathogens by chance, due to a compromised immune system potentially related to inbreeding, or as a result of specific environmental conditions influencing pathogen loads in certain populations. The geographic distribution of the outlier desmans may provide a clue, as four of them belonged to the same hydrological unit, Endrinal, so that 40% of the desmans in this subpopulation would have an elevated or altered pathogen load. The Endrinal river supports a significant livestock density (personal observation of the authors), which could be behind the unique pathogen load found in the desmans of this river. However, additional sampling and detailed environmental analyses in this and other hydrological units is needed to better understand the origin of these pathogens and the factors driving the altered pathogen loads in some desmans and populations.

Future studies of Iberian desman pathogens should include different populations across the species’ range and throughout the year to understand population health trends and to prioritize conservation efforts to the most vulnerable populations. Integrating the analysis of genetic factors, such as inbreeding and mutational load, will be crucial for understanding their influence on pathogen load and susceptibility. Furthermore, epidemiological studies should be a priority to understand how these pathogens are transmitted, for example through water or through contact with other wildlife species, to identify potential transmission hotspots, such as excessive livestock densities in mountainous areas or wastewater discharge points near human settlements, and to assess their potential impact on the health and viability of Iberian desman populations.

### Method challenges and prospects

This study represents a significant advance in pathogen detection methods for endangered species by providing a novel, effective, and non-invasive metagenomic approach for monitoring microbial diseases. To demonstrate the effectiveness of the method, we used a dataset of 136 bacterial reference genomes for mapping. We focused on bacteria because of their adequate representation of complete genomes in databases, as well as their reduced genome sizes, which allowed us to test a wide variety of parameters and analysis conditions with reasonable computational effort. Viral and eukaryotic pathogens, on the other hand, present unique challenges for metagenomic detection, which were beyond the scope of this study. Future research should address these challenges and optimize tools for detecting such pathogens, as they can also have a significant impact on the overall health and viability of wildlife species (Nunn & Altizer, 2005; Pedersen et al., 2007).

The dependence on reference genome databases is a major limitation of mapping methods, particularly for less studied species such as the Iberian desman. Our results showed that many pathogenic bacterial species present in genome databases (19 out of 136 analyzed) can be found in the Iberian desman. Most of these bacteria also infect humans, suggesting a wide host range for these pathogens (Shaw et al., 2020). However, since bacterial pathogens specific to the Iberian desman have not yet been sequenced, only closely related species present in databases can be detected using this bioinformatics method. Eukaryotic parasite species, especially those with complex life cycles, tend to be more host-specific (Poulin, 2006), so it remains to be seen whether genomes currently present in genome databases, usually isolated from humans, will be useful for detecting eukaryotic pathogens in the Iberian desman or other endangered mammals. In the long term, the generation of complete reference genomes for a wide range of relevant pathogenic and parasitic species will be key to advancing the detection and assessment of pathogens in wildlife, especially in critically endangered species.

Accurate identification of pathogens is not only critical in conservation biology, but also an integral part of the One Health approach, which emphasizes the important interrelationships that exist between human, animal, and environmental health (Destoumieux-Garzon et al., 2018; White & Razgour, 2020). By leveraging advances in high-throughput sequencing and metagenomic technologies, this study provides a reliable and non-invasive method to monitor pathogens that may pose risks, not only to wildlife and endangered species, but also to humans. This underscores the importance of further research in this area, particularly to improve genomic tools and the representation of reference genomes of wildlife pathogens and parasites in databases, thereby contributing to the broader goal of protecting endangered species and maintaining ecosystem health.

## Data Availability

Upon acceptance for publication in a peer-reviewed journal, data will be deposited in a digital repository.

## Supplementary Material

Additional data and figures are available in Supplementary Material.

## Acknowledgements

This work was funded by research grant TED2021-130149B-I00 of MCIN/AEI/10.13039/501100011033 and the European Union NextGenerationEU/PRTR. Fieldwork was supported by the Ministry of the Environment through the Biodiversity Foundation, the Duero River Basin Authority, and the Autonomous Government of Castilla y León through the Patrimonio Natural Foundation. We also thank people of Biosfera who helped in fieldwork, especially Sergi Munné Prat, Alejandro González Ibáñez, and Jose María Valle Artaza.

