## Supplementary Information for "Development of metagenomic methods for non-invasive health monitoring of endangered species: Unveiling hidden microbial threats in fecal samples"

**Table S1.** Samples used in this study, hydrological unit or population of origin of the sample, and number of exogenous reads after filtering out endogenous reads.

| Sample number | Sample | Hydrological unit | Exogenous reads |
|---|---|---|---|
| 1 | BC2042 | Aravalle | 61,736,980 |
| 2 | BC2045 | Endrinal | 50,124,430 |
| 3 | BC2280 | Adaja | 59,649,976 |
| 4 | BC2360 | Adaja | 56,053,206 |
| 5 | BC2378 | Endrinal | 48,532,104 |
| 6 | BC2381 | Endrinal | 60,256,352 |
| 7 | BC2408 | Aravalle | 77,905,478 |
| 8 | BC2416 | Adaja | 66,786,794 |
| 9 | BC2765 | Becedillas | 47,017,792 |
| 10 | BC2856 | Becedillas | 81,444,940 |
| 11 | BC2865 | Endrinal | 30,409,082 |
| 12 | BC2876 | Aravalle | 65,926,618 |
| 13 | BC2934 | Adaja | 72,800,924 |
| 14 | BC2989 | Becedillas | 63,910,586 |
| 15 | BC3015 | Aravalle | 41,132,864 |
| 16 | BC3267 | Aravalle | 75,036,328 |
| 17 | BC3343 | Aravalle | 41,438,942 |
| 18 | BC3345 | Endrinal | 53,654,356 |
| 19 | BC3876 | Endrinal | 78,895,038 |
| 20 | BC3895 | Endrinal | 66,061,256 |
| 21 | BC3899 | Endrinal | 29,719,132 |
| 22 | BC3903 | Endrinal | 43,476,990 |
| 23 | BC3906 | Endrinal | 56,744,902 |

**Table S2.** Species of the genus *Yersinia* according to NCBI Taxonomy, along with the size of the reference genome used. Pathogenicity is indicated with an X.

| Species | Genome size (bp) | Source | Pathogenicity |
|---|---|---|---|
| *Yersinia aldovae* | 4,471,090 | NCBI Taxonomy | |
| *Yersinia aleksiciae* | 4,526,044 | NCBI Taxonomy | |
| *Yersinia alsatica* | 4,901,396 | NCBI Taxonomy | |
| *Yersinia artesiana* | 4,520,664 | NCBI Taxonomy | |
| *Yersinia bercovieri* | 4,428,721 | NCBI Taxonomy | |
| *Yersinia canariae* | 4,710,154 | NCBI Taxonomy | |
| *Yersinia enterocolitica* | 4,548,822 | NCBI Taxonomy | X |
| *Yersinia entomophaga* | 4,275,406 | NCBI Taxonomy | |
| *Yersinia frederiksenii* | 4,941,872 | NCBI Taxonomy | |
| *Yersinia hibernica* | 4,803,435 | NCBI Taxonomy | |
| *Yersinia intermedia* | 4,928,910 | NCBI Taxonomy | |
| *Yersinia kristensenii* | 4,733,508 | NCBI Taxonomy | |
| *Yersinia massiliensis* | 5,050,276 | NCBI Taxonomy | |
| *Yersinia mollaretii* | 4,603,534 | NCBI Taxonomy | |
| *Yersinia nurmii* | 4,143,134 | NCBI Taxonomy | |
| *Yersinia pekkanenii* | 5,046,986 | NCBI Taxonomy | |
| *Yersinia pestis* | 4,658,411 | NCBI Taxonomy | X |
| *Yersinia proxima* | 4,616,898 | NCBI Taxonomy | |
| *Yersinia pseudotuberculosis* | 4,839,430 | NCBI Taxonomy | X |
| *Yersinia rochesterensis* | 4,448,883 | NCBI Taxonomy | |
| *Yersinia rohdei* | 4,372,253 | NCBI Taxonomy | |
| *Yersinia ruckeri* | 3,894,226 | NCBI Taxonomy | X |
| *Yersinia similis* | 4,964,409 | NCBI Taxonomy | |
| *Yersinia thracica* | 4,420.333 | NCBI Taxonomy | |
| *Yersinia vastinensis* | 4,540,093 | NCBI Taxonomy | |
| *Yersinia wautersii* | 4,858,848 | NCBI Taxonomy | |

**Table S3.** Species of pathogenic bacteria analyzed and the size of their reference genomes.

| Species | Synonym | Genome size (bp) |
|---|---|---|
| *Acinetobacter baumannii* | | 3,980,230 |
| *Actinobacillus lignieresii* | | 2,236,520 |
| *Aeromonas hydrophila* | | 4,733,702 |
| *Aeromonas salmonicida* | | 4,954,811 |
| *Aeromonas sobria* | | 4,683,669 |
| *Aeromonas veronii* | | 4,561,870 |
| *Anaplasma marginale* | | 1,202,435 |
| *Anaplasma phagocytophilum* | | 1,481,598 |
| *Bacillus anthracis* | | 5,503,926 |
| *Bacillus cereus* | | 5,836,971 |
| *Bartonella bacilliformis* | | 1,411,655 |
| *Bartonella henselae* | | 1,905,383 |
| *Bartonella quintana* | | 1,588,683 |
| *Bordetella pertussis* | | 4,088,701 |
| *Borrelia afzelii* | | 906,136 |
| *Borrelia burgdorferi* | | 1,321,434 |
| *Borrelia garinii* | | 905,692 |
| *Brucella abortus* | | 3,278,307 |
| *Brucella melitensis* | | 3,294,931 |
| *Brucella microti* | | 3,337,369 |
| *Brucella neotomae* | | 3,329,628 |
| *Brucella ovis* | | 3,275,590 |
| *Brucella suis* | | 3,315,175 |
| *Burkholderia cepacia* | | 8,366,868 |
| *Burkholderia pseudomallei* | | 7,085,397 |
| *Campylobacter coli* | | 1,716,536 |
| *Campylobacter fetus* | | 1,804,582 |
| *Campylobacter jejuni* | | 1,641,481 |
| *Chlamydia psittaci* | | 1,179,220 |
| *Chlamydia trachomatis* | | 1,042,519 |
| *Citrobacter freundii* | | 5,171,093 |
| *Clostridioides difficile* | *Clostridium difficile* | 4,095,894 |
| *Clostridium botulinum* | | 3,903,260 |
| *Clostridium chauvoei* | | 2,889,569 |
| *Clostridium histolyticum* | | 2,740,791 |
| *Clostridium novyi* | | 2,499,078 |
| *Clostridium perfringens* | | 3,275,424 |
| *Clostridium septicum* | | 3,404,718 |
| *Clostridium tetani* | | 2,873,333 |
| *Corynebacterium diphtheriae* | | 2,406,896 |
| *Corynebacterium striatum* | | 2,904,831 |
| *Coxiella burnetii* | | 2,032,807 |
| *Edwardsiella ictaluri* | | 3,844,237 |
| *Edwardsiella piscicida* | | 3,819,771 |
| *Edwardsiella tarda* | | 3,720,168 |
| *Elizabethkingia anophelis* | | 4,058,311 |

| | | |
|---|---|---|
| *Enterobacter asburiae* | | 4,768,325 |
| *Enterobacter bugandensis* | | 4,734,039 |
| *Enterobacter cancerogenus* | | 4,736,684 |
| *Enterobacter chengduensis* | | 5,218,125 |
| *Enterobacter chuandaensis* | | 4,634,324 |
| *Enterobacter cloacae* | | 5,023,439 |
| *Enterobacter hormaechei* | | 4,855,498 |
| *Enterobacter kobei* | | 4,773,801 |
| *Enterobacter ludwigii* | | 4,952,770 |
| *Enterobacter mori* | | 4,844,012 |
| *Enterobacter oligotrophicus* | | 4,476,585 |
| *Enterobacter roggenkampii* | | 4,899,997 |
| *Enterobacter sichuanensis* | | 4,711,389 |
| *Enterobacter soli* | | 5,012,132 |
| *Enterococcus faecalis* | | 2,870,381 |
| *Enterococcus faecium* | | 2,919,198 |
| *Enterococcus hirae* | | 2,845,651 |
| *Escherichia coli* | | 5,594,605 |
| *Flavobacterium psychrophilum* | | 2,830,557 |
| *Francisella tularensis* | | 1,870,206 |
| *Fusobacterium necrophorum* | | 2,288,480 |
| *Haemophilus influenzae* | | 1,846,259 |
| *Helicobacter pylori* | | 1,624,458 |
| *Klebsiella oxytoca* | | 5,879,076 |
| *Klebsiella pneumoniae* | | 5,682,322 |
| *Kluyvera intermedia* | | 4,938,529 |
| *Kosakonia oryzendophytica* | | 5,132,919 |
| *Kosakonia oryziphila* | | 4,814,900 |
| *Legionella pneumophila* | | 3,504,074 |
| *Leptospira interrogans* | | 4,630,763 |
| *Listeria innocua* | | 2,922,148 |
| *Listeria ivanovii* | | 2,919,548 |
| *Listeria monocytogenes* | | 2,944,528 |
| *Listeria seeligeri* | | 2,797,636 |
| *Mannheimia haemolytica* | | 2,757,078 |
| *Mesomycoplasma conjunctivae* | *Mycoplasma conjunctivae* | 846,214 |
| *Mesomycoplasma hyopneumoniae* | *Mycoplasma hyopneumoniae* | 921,093 |
| *Morganella morganii* | | 3,906,921 |
| *Mycobacterium bovis* | *M. tuberculosis variant bovis* | 4,411,814 |
| *Mycobacterium caprae* | *M. tuberculosis variant caprae* | 4,324,961 |
| *Mycobacterium microti* | *M. tuberculosis variant microti* | 4,369,915 |
| *Mycobacterium tuberculosis* | | 4,411,532 |
| *Mycoplasma mycoides* | | 1,084,586 |
| *Mycoplasmoides pneumoniae* | *Mycoplasma pneumoniae* | 823,017 |
| *Mycoplasmopsis agalactiae* | *Mycoplasma agalactiae* | 916,806 |
| *Neisseria gonorrhoeae* | | 2,171,755 |
| *Neisseria lactamica* | | 2,200,224 |
| *Neisseria meningitidis* | | 2,157,444 |
| *Neisseria polysaccharea* | | 2,029,584 |

| | |
|---|---|
| *Pasteurella multocida* | 2,335,516 |
| *Photobacterium damselae* | 4,427,003 |
| *Phytobacter massiliensis* | 5,003,540 |
| *Pluralibacter gergoviae* | 5,408,082 |
| *Providencia alcalifaciens* | 3,990,106 |
| *Pseudomonas aeruginosa* | 6,264,404 |
| *Pseudomonas putida* | 6,156,701 |
| *Rickettsia conorii* | 1,268,755 |
| *Rickettsia monacensis* | 1,353,450 |
| *Rickettsia prowazekii* | 1,109,804 |
| *Rickettsia rickettsii* | 1,257,710 |
| *Rickettsia typhi* | 1,112,372 |
| *Salmonella bongori* | 4,773,537 |
| *Salmonella enterica* | 4,951,383 |
| *Serratia marcescens* | 5,238,537 |
| *Shewanella algae* | 4,990,025 |
| *Shigella dysenteriae* | 5,192,674 |
| *Shigella flexneri* | 4,828,820 |
| *Shigella sonnei* | 4,762,774 |
| *Staphylococcus aureus* | 2,821,361 |
| *Staphylococcus pseudintermedius* | 2,629,596 |
| *Stenotrophomonas maltophilia* | 4,481,118 |
| *Streptobacillus moniliformis* | 1,673,280 |
| *Streptococcus agalactiae* | 2,079,123 |
| *Streptococcus equi* | 2,154,778 |
| *Streptococcus iniae* | 2,078,360 |
| *Streptococcus mutans* | 2,028,032 |
| *Streptococcus pneumoniae* | 2,154,362 |
| *Streptococcus pyogenes* | 1,746,380 |
| *Streptococcus suis* | 2,170,808 |
| *Treponema pallidum* | 1,139,330 |
| *Vibrio cholerae* | 4,138,412 |
| *Vibrio fluvialis* | 4,827,733 |
| *Vibrio harveyi* | 5,869,008 |
| *Vibrio metschnikovii* | 3,831,334 |
| *Vibrio parahaemolyticus* | 5,165,770 |
| *Vibrio vulnificus* | 5,117,890 |
| *Yersinia enterocolitica* | 4,548,822 |
| *Yersinia pestis* | 4,658,411 |
| *Yersinia pseudotuberculosis* | 4,839,430 |
| *Yersinia ruckeri* | 3,894,226 |

**Table S4.** Number of reads and breadth of coverage (%) obtained from aligning reads with *Yersinia* species genomes using Bowtie2 in end-to-end mode with default parameters.

| Number of Reads | BC2042 | BC2045 | BC2280 | BC2360 | BC2378 | BC2381 | BC2408 | BC2416 | BC2765 | BC2856 | BC2865 | BC2876 | BC2934 | BC2989 | BC3015 | BC3267 | BC3343 | BC3345 | BC3876 | BC3895 | BC3899 | BC3903 | BC3906 |
|---|---|---|---|---|---|---|---|---|---|---|---|---|---|---|---|---|---|---|---|---|---|---|---|
| *Y. aldovae* | 86 | 100 | 30 | 22 | 104 | 262 | 288 | 6 | 230 | 10 | 44 | 468 | 308 | 164 | 224 | 126 | 58 | 13790 | 338 | 32 | 272 | 12378 | 76 |
| *Y. aleksiciae* | 278 | 4048 | 1470 | 1120 | 1718 | 7118 | 2900 | 60 | 2498 | 46 | 1502 | 2788 | 12538 | 1792 | 2490 | 92 | 266 | 20016 | 6566 | 148 | 1462 | 624 | 384 |
| *Y. alsatica* | 50 | 90 | 42 | 20 | 102 | 70 | 228 | 2 | 322 | 8 | 44 | 558 | 332 | 88 | 1188 | 10 | 24 | 5636 | 404 | 2 | 250 | 406 | 92 |
| *Y. artesiana* | 298 | 4600 | 1664 | 1256 | 1970 | 7798 | 3200 | 68 | 2866 | 56 | 1676 | 3090 | 14842 | 1958 | 2822 | 98 | 286 | 19082 | 7702 | 150 | 1610 | 868 | 426 |
| *Y. bercovieri* | 282 | 4308 | 1542 | 1204 | 1826 | 7528 | 3110 | 62 | 2562 | 50 | 1546 | 2826 | 13418 | 1782 | 2610 | 98 | 268 | 19044 | 6850 | 148 | 1516 | 588 | 364 |
| *Y. canariae* | 24 | 34 | 70 | 10 | 86 | 76 | 220 | 2 | 252 | 4 | 30 | 492 | 110 | 56 | 244 | 8 | 30 | 5700 | 360 | 6 | 216 | 390 | 84 |
| *Y. enterocolitica* | 38 | 84 | 82 | 20 | 128 | 108 | 256 | 4 | 336 | 6 | 64 | 604 | 1364 | 40 | 316 | 14 | 34 | 6352 | 456 | 8 | 326 | 976 | 116 |
| *Y. entomophaga* | 260 | 110 | 50 | 18 | 164 | 446 | 188 | 0 | 132 | 4 | 50 | 202 | 446 | 88 | 212 | 112 | 28 | 2494 | 16966 | 0 | 100 | 122 | 38 |
| *Y. frederiksenii* | 36 | 126 | 92 | 32 | 122 | 138 | 262 | 12 | 334 | 6 | 32 | 514 | 454 | 92 | 332 | 18 | 22 | 6310 | 442 | 6 | 266 | 386 | 88 |
| *Y. hibernica* | 24 | 284 | 50 | 30 | 86 | 68 | 186 | 2 | 226 | 4 | 36 | 378 | 300 | 58 | 208 | 12 | 32 | 5876 | 292 | 4 | 196 | 324 | 70 |
| *Y. intermedia* | 832 | 214 | 148 | 164 | 4724 | 1166 | 9908 | 14 | 22870 | 24 | 562 | 32400 | 314 | 120 | 10780 | 76 | 878 | 29610 | 9128 | 456 | 17292 | 904 | 7162 |
| *Y. kristensenii* | 40 | 44 | 20 | 12 | 64 | 80 | 218 | 6 | 234 | 4 | 24 | 428 | 138 | 44 | 206 | 72 | 26 | 8670 | 410 | 2 | 208 | 414 | 84 |
| *Y. massiliensis* | 286 | 4582 | 1706 | 1318 | 1944 | 7540 | 3188 | 62 | 2588 | 54 | 1586 | 2872 | 14250 | 1856 | 2702 | 100 | 278 | 19978 | 7410 | 156 | 1518 | 508 | 368 |
| *Y. mollaretii* | 28 | 64 | 30 | 8 | 92 | 112 | 222 | 6 | 278 | 6 | 24 | 488 | 210 | 54 | 218 | 10 | 32 | 4878 | 330 | 2 | 234 | 326 | 94 |
| *Y. nurmii* | 304 | 5234 | 1842 | 1368 | 1916 | 7864 | 2946 | 62 | 2550 | 50 | 1540 | 2546 | 15304 | 2078 | 2564 | 102 | 264 | 14782 | 9348 | 146 | 1478 | 316 | 334 |
| *Y. pekkanenii* | 300 | 4324 | 1544 | 1178 | 1810 | 7500 | 2862 | 64 | 2538 | 50 | 1632 | 2774 | 13532 | 2006 | 2584 | 102 | 256 | 19672 | 6938 | 142 | 1494 | 760 | 374 |
| *Y. pestis* | 28 | 74 | 38 | 14 | 66 | 58 | 174 | 4 | 122 | 4 | 22 | 184 | 226 | 44 | 126 | 6 | 24 | 4816 | 204 | 0 | 138 | 200 | 42 |
| *Y. proxima* | 290 | 4754 | 1610 | 1284 | 1894 | 7750 | 3192 | 64 | 2668 | 56 | 1592 | 3020 | 14136 | 1908 | 3172 | 100 | 282 | 19378 | 7222 | 154 | 1606 | 990 | 412 |
| *Y. pseudotuberculosis* | 26 | 62 | 38 | 20 | 76 | 78 | 132 | 6 | 122 | 4 | 32 | 202 | 1020 | 40 | 122 | 6 | 24 | 6172 | 306 | 0 | 146 | 202 | 46 |
| *Y. rochesterensis* | 26 | 86 | 50 | 28 | 90 | 106 | 216 | 6 | 266 | 2 | 26 | 454 | 246 | 34 | 200 | 10 | 22 | 5282 | 318 | 2 | 222 | 386 | 82 |
| *Y. rohdei* | 24 | 102 | 36 | 30 | 92 | 74 | 184 | 6 | 316 | 4 | 28 | 460 | 370 | 48 | 248 | 12 | 28 | 4880 | 380 | 6 | 244 | 412 | 90 |
| *Y. ruckeri* | 92 | 110 | 26 | 10 | 100 | 74 | 154 | 2 | 78 | 4 | 20 | 214 | 224 | 62 | 154 | 20 | 78 | 3998 | 1650 | 2 | 90 | 138 | 40 |
| *Y. similis* | 34 | 48 | 12 | 4 | 66 | 44 | 218 | 4 | 146 | 4 | 28 | 238 | 94 | 50 | 132 | 4 | 16 | 4686 | 192 | 0 | 128 | 218 | 38 |
| *Y. thracica* | 276 | 3784 | 1406 | 1086 | 1696 | 7498 | 2876 | 60 | 2360 | 42 | 1610 | 2752 | 12316 | 1786 | 2502 | 94 | 268 | 19714 | 6402 | 138 | 1578 | 660 | 352 |
| *Y. vastinensis* | 308 | 5076 | 1772 | 1332 | 1978 | 7840 | 3090 | 58 | 2810 | 56 | 1778 | 2986 | 15352 | 2104 | 2724 | 96 | 284 | 18480 | 7754 | 156 | 1608 | 532 | 382 |
| *Y. wautersii* | 182 | 2970 | 1040 | 898 | 1308 | 5658 | 1754 | 48 | 2142 | 38 | 932 | 1508 | 9926 | 1258 | 1452 | 62 | 150 | 13868 | 4812 | 82 | 992 | 418 | 218 |

| Breadth of Coverage | BC2042 | BC2045 | BC2280 | BC2360 | BC2378 | BC2381 | BC2408 | BC2416 | BC2765 | BC2856 | BC2865 | BC2876 | BC2934 | BC2989 | BC3015 | BC3267 | BC3343 | BC3345 | BC3876 | BC3895 | BC3899 | BC3903 | BC3906 |
|---|---|---|---|---|---|---|---|---|---|---|---|---|---|---|---|---|---|---|---|---|---|---|---|
| *Y. aldovae* | 0.19 | 0.1 | 0.04 | 0.05 | 0.16 | 0.17 | 0.52 | 0.01 | 0.37 | 0.02 | 0.08 | 0.77 | 0.26 | 0.34 | 0.37 | 0.27 | 0.13 | 1.5 | 0.49 | 0.08 | 0.44 | 19.03 | 0.16 |
| *Y. aleksiciae* | 0.13 | 0.16 | 0.14 | 0.11 | 0.25 | 0.22 | 0.48 | 0.06 | 0.52 | 0.06 | 0.13 | 0.85 | 0.26 | 0.18 | 0.45 | 0.09 | 0.15 | 0.96 | 0.5 | 0.08 | 0.47 | 0.66 | 0.26 |
| *Y. alsatica* | 0.1 | 0.07 | 0.05 | 0.03 | 0.16 | 0.09 | 0.34 | 0 | 0.46 | 0.01 | 0.03 | 0.87 | 0.14 | 0.1 | 2.25 | 0.02 | 0.05 | 0.88 | 0.42 | 0 | 0.38 | 0.56 | 0.18 |
| *Y. artesiana* | 0.17 | 0.19 | 0.12 | 0.12 | 0.28 | 0.41 | 0.5 | 0.07 | 0.6 | 0.06 | 0.14 | 0.88 | 0.54 | 0.17 | 0.63 | 0.11 | 0.15 | 1.12 | 0.76 | 0.09 | 0.52 | 1.01 | 0.28 |
| *Y. bercovieri* | 0.14 | 0.15 | 0.13 | 0.12 | 0.25 | 0.22 | 0.51 | 0.06 | 0.56 | 0.06 | 0.13 | 0.82 | 0.22 | 0.17 | 0.46 | 0.1 | 0.13 | 0.95 | 0.49 | 0.08 | 0.49 | 0.61 | 0.26 |
| *Y. canariae* | 0.05 | 0.05 | 0.04 | 0.01 | 0.13 | 0.12 | 0.32 | 0 | 0.4 | 0.01 | 0.04 | 0.78 | 0.12 | 0.06 | 0.39 | 0.02 | 0.07 | 0.94 | 0.5 | 0.01 | 0.36 | 0.55 | 0.17 |
| *Y. enterocolitica* | 0.08 | 0.07 | 0.12 | 0.03 | 0.19 | 0.14 | 0.39 | 0.01 | 0.52 | 0.02 | 0.07 | 0.91 | 2.34 | 0.06 | 0.53 | 0.03 | 0.08 | 1.19 | 0.48 | 0.02 | 0.51 | 1.54 | 0.23 |
| *Y. entomophaga* | 0.67 | 0.09 | 0.06 | 0.02 | 0.28 | 1 | 0.24 | 0 | 0.17 | 0.01 | 0.05 | 0.32 | 0.14 | 0.11 | 0.34 | 0.27 | 0.06 | 0.4 | 30.63 | 0 | 0.14 | 0.2 | 0.08 |
| *Y. frederiksenii* | 0.06 | 0.07 | 0.06 | 0.04 | 0.17 | 0.15 | 0.34 | 0.02 | 0.45 | 0.01 | 0.03 | 0.73 | 0.15 | 0.1 | 0.46 | 0.04 | 0.04 | 0.96 | 0.37 | 0.01 | 0.4 | 0.52 | 0.16 |
| *Y. hibernica* | 0.04 | 0.13 | 0.04 | 0.05 | 0.15 | 0.1 | 0.28 | 0 | 0.34 | 0.01 | 0.05 | 0.57 | 0.19 | 0.07 | 0.33 | 0.03 | 0.07 | 0.8 | 0.35 | 0.01 | 0.31 | 0.46 | 0.14 |
| *Y. intermedia* | 1.8 | 0.34 | 0.27 | 0.4 | 8.47 | 2.44 | 18.36 | 0.02 | 34.52 | 0.05 | 0.99 | 45.38 | 0.39 | 0.22 | 18.78 | 0.16 | 1.79 | 37.85 | 15.81 | 1.03 | 27.54 | 1.47 | 13.69 |
| *Y. kristensenii* | 0.08 | 0.04 | 0.03 | 0.03 | 0.1 | 0.13 | 0.31 | 0.01 | 0.37 | 0.01 | 0.04 | 0.72 | 0.13 | 0.08 | 0.35 | 0.05 | 0.06 | 0.8 | 0.47 | 0 | 0.33 | 0.57 | 0.18 |
| *Y. massiliensis* | 0.12 | 0.13 | 0.1 | 0.09 | 0.18 | 0.15 | 0.32 | 0.05 | 0.33 | 0.05 | 0.11 | 0.56 | 0.18 | 0.13 | 0.31 | 0.09 | 0.12 | 0.73 | 0.36 | 0.06 | 0.31 | 0.43 | 0.19 |
| *Y. mollaretii* | 0.06 | 0.07 | 0.06 | 0.02 | 0.14 | 0.12 | 0.36 | 0.01 | 0.45 | 0.01 | 0.04 | 0.82 | 0.13 | 0.09 | 0.37 | 0.02 | 0.07 | 0.87 | 0.38 | 0 | 0.4 | 0.48 | 0.2 |
| *Y. nurmii* | 0.19 | 0.18 | 0.15 | 0.11 | 0.2 | 0.28 | 0.27 | 0.06 | 0.23 | 0.05 | 0.13 | 0.35 | 0.2 | 0.2 | 0.29 | 0.12 | 0.13 | 0.48 | 3.35 | 0.07 | 0.22 | 0.28 | 0.15 |
| *Y. pekkanenii* | 0.16 | 0.13 | 0.1 | 0.09 | 0.19 | 0.18 | 0.41 | 0.06 | 0.49 | 0.05 | 0.11 | 0.84 | 0.22 | 0.15 | 0.4 | 0.08 | 0.11 | 0.98 | 0.4 | 0.07 | 0.44 | 0.7 | 0.25 |
| *Y. pestis* | 0.06 | 0.06 | 0.03 | 0.02 | 0.09 | 0.08 | 0.21 | 0.01 | 0.18 | 0.01 | 0.03 | 0.3 | 0.1 | 0.07 | 0.18 | 0.01 | 0.05 | 0.55 | 0.21 | 0 | 0.19 | 0.3 | 0.08 |
| *Y. proxima* | 0.15 | 0.16 | 0.12 | 0.15 | 0.24 | 0.21 | 0.43 | 0.06 | 0.53 | 0.06 | 0.14 | 0.88 | 0.48 | 0.17 | 1.23 | 0.1 | 0.15 | 1.09 | 0.55 | 0.07 | 0.47 | 1.23 | 0.27 |
| *Y. pseudotuberculosis* | 0.05 | 0.05 | 0.03 | 0.03 | 0.1 | 0.09 | 0.18 | 0.01 | 0.18 | 0.01 | 0.04 | 0.31 | 0.71 | 0.06 | 0.18 | 0.01 | 0.05 | 0.57 | 0.23 | 0 | 0.19 | 0.29 | 0.08 |
| *Y. rochesterensis* | 0.06 | 0.06 | 0.04 | 0.04 | 0.13 | 0.17 | 0.35 | 0.01 | 0.42 | 0 | 0.04 | 0.78 | 0.17 | 0.06 | 0.33 | 0.02 | 0.05 | 0.88 | 0.43 | 0 | 0.39 | 0.61 | 0.18 |
| *Y. rohdei* | 0.05 | 0.07 | 0.04 | 0.04 | 0.13 | 0.12 | 0.31 | 0.01 | 0.45 | 0.01 | 0.04 | 0.75 | 0.16 | 0.08 | 0.41 | 0.02 | 0.07 | 0.93 | 0.43 | 0.02 | 0.41 | 0.7 | 0.19 |
| *Y. ruckeri* | 0.25 | 0.08 | 0.03 | 0.01 | 0.17 | 0.14 | 0.21 | 0 | 0.13 | 0.01 | 0.04 | 0.41 | 0.15 | 0.11 | 0.26 | 0.05 | 0.21 | 0.59 | 3.64 | 0 | 0.17 | 0.23 | 0.09 |
| *Y. similis* | 0.07 | 0.04 | 0.02 | 0.01 | 0.1 | 0.07 | 0.26 | 0.01 | 0.2 | 0.01 | 0.04 | 0.32 | 0.08 | 0.05 | 0.18 | 0.01 | 0.04 | 0.54 | 0.2 | 0 | 0.19 | 0.28 | 0.07 |
| *Y. thracica* | 0.15 | 0.14 | 0.11 | 0.11 | 0.22 | 0.24 | 0.47 | 0.06 | 0.54 | 0.05 | 0.11 | 0.83 | 0.2 | 0.18 | 0.49 | 0.09 | 0.16 | 1.03 | 0.56 | 0.08 | 0.58 | 0.7 | 0.26 |
| *Y. vastinensis* | 0.15 | 0.15 | 0.14 | 0.12 | 0.26 | 0.23 | 0.44 | 0.05 | 0.57 | 0.06 | 0.14 | 0.84 | 0.22 | 0.19 | 0.47 | 0.1 | 0.15 | 0.97 | 0.53 | 0.07 | 0.46 | 0.56 | 0.26 |
| *Y. wautersii* | 0.13 | 0.15 | 0.12 | 0.12 | 0.21 | 0.28 | 0.36 | 0.04 | 0.44 | 0.04 | 0.1 | 0.42 | 0.2 | 0.14 | 0.27 | 0.06 | 0.1 | 0.7 | 0.44 | 0.05 | 0.27 | 0.39 | 0.15 |

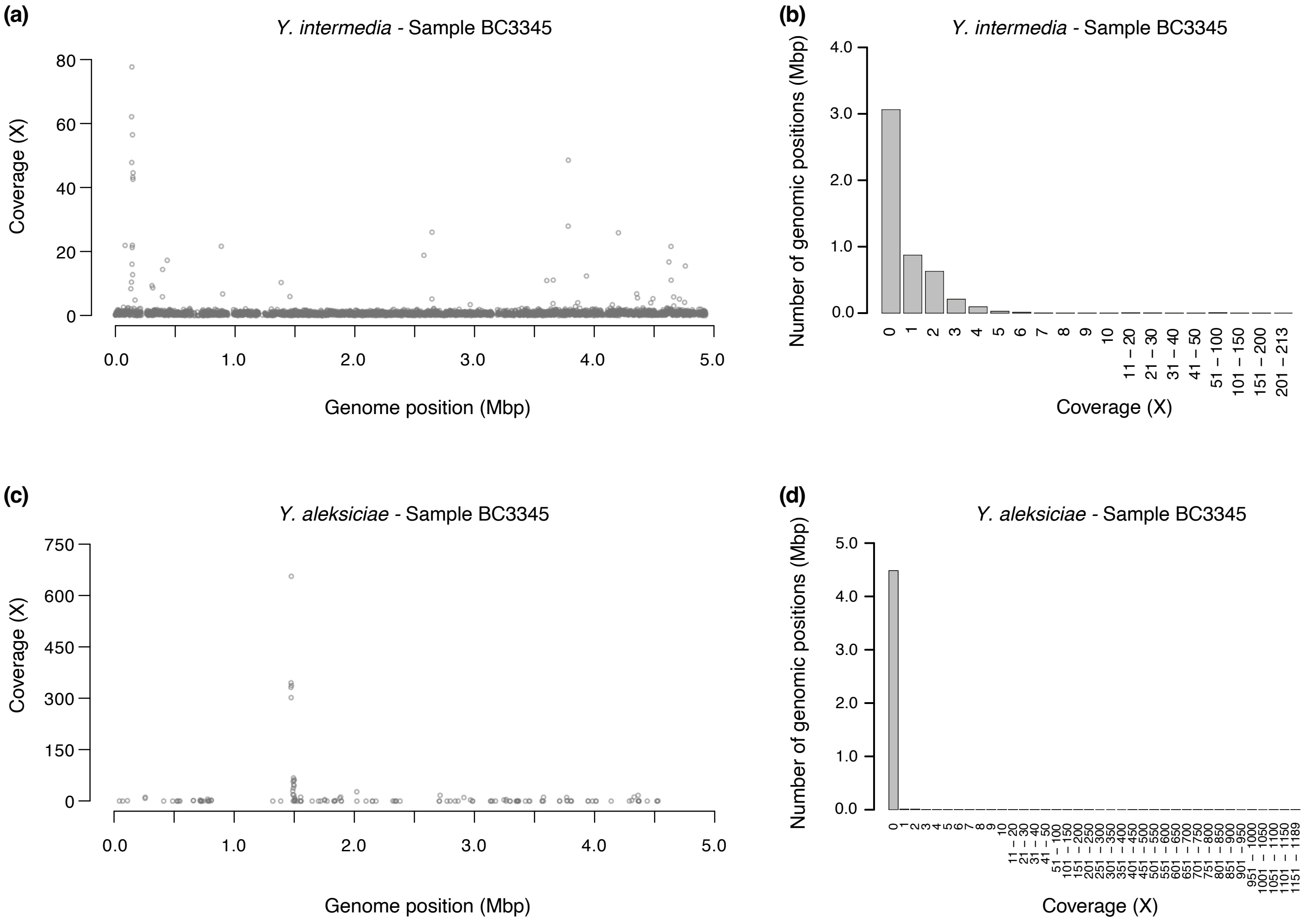


**Figure S1.** Comparison of depth of coverage across the reference genomes of *Y. intermedia* and *Y. aleksiciae* and the corresponding depth of coverage barplots for sample BC3345. (a) Coverage across the reference genome of *Y. intermedia* in 4,000 windows and (b) coverage barplot, in which 29,610 reads were assigned, resulting in a mean depth coverage of 0.90X and a breadth of coverage of 37.85%. (c) Coverage across the reference genome of *Y. aleksiciae* and (d) coverage barplot, in which 20,016 reads were assigned, resulting in a mean depth coverage of 0.66X and a breadth of coverage of 0.96%. In (a) and (c), windows with zero coverage are not plotted. Note the different coverage scales in (a) and (c).

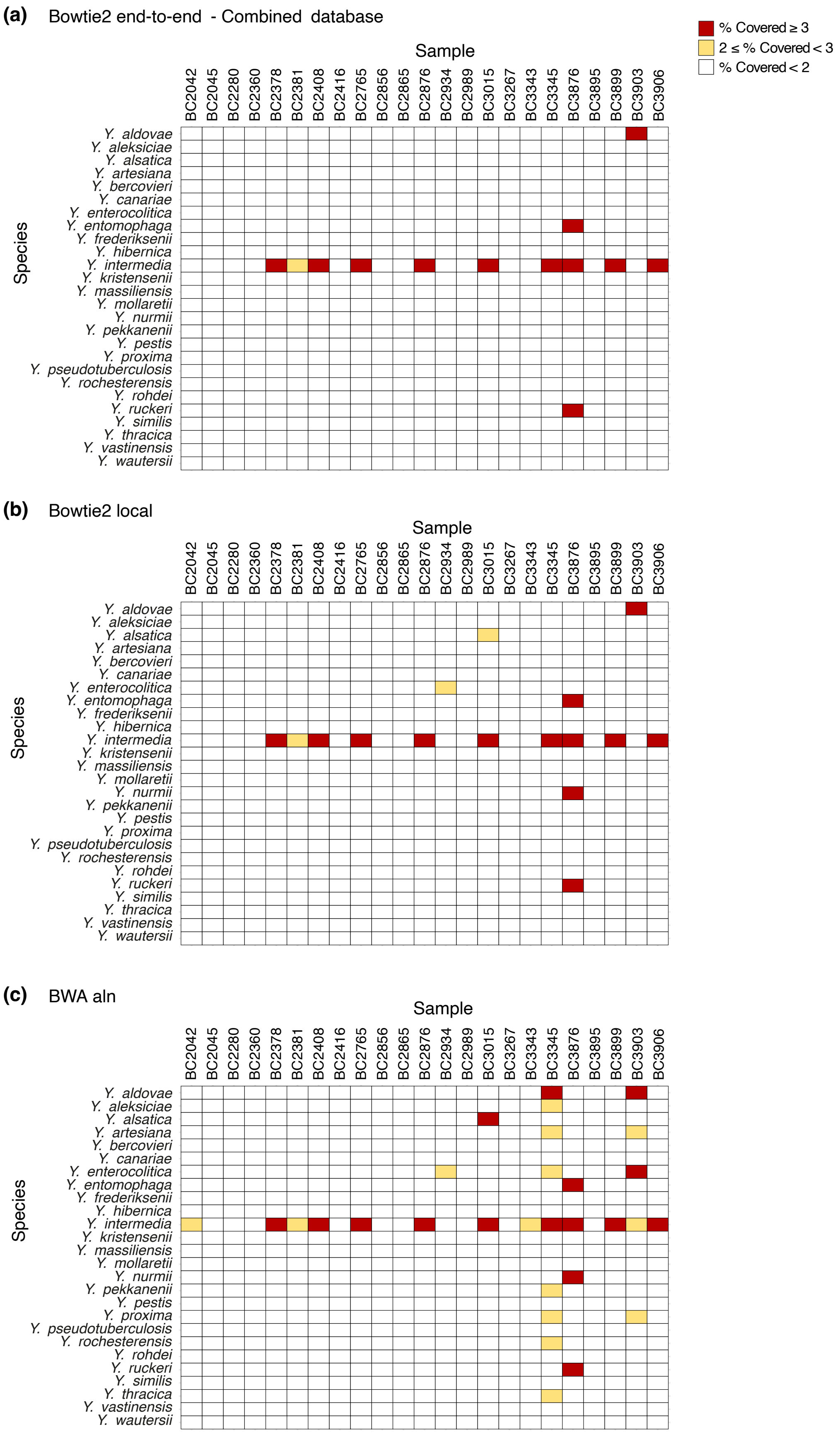


**Figure S2.** Identification of *Yersinia* species in 23 samples of Iberian desman using different approaches. (a) Identification with a combined database for all species using Bowtie2 in end-to-end mode. (b) Identification with single-species databases using Bowtie2 in default local mode. (c) Identification with single-species databases using BWA-aln with default parameters.

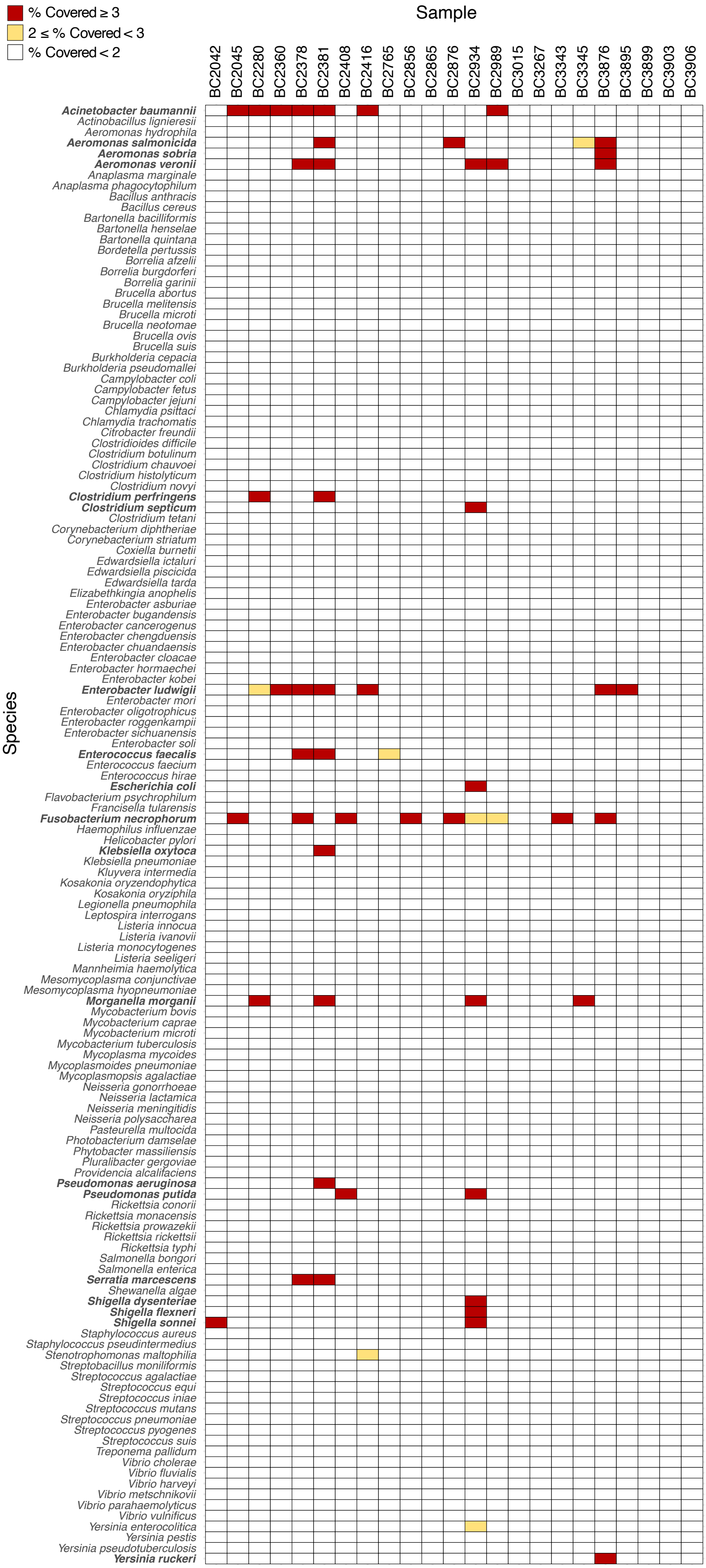


**Figure S3**. Identification of 136 bacterial species in 23 samples of Iberian desman using a combined database. Species were identified according to the percentage of the genome covered by mapped reads. Species with coverage greater than 3% are shown in bold for clarity.

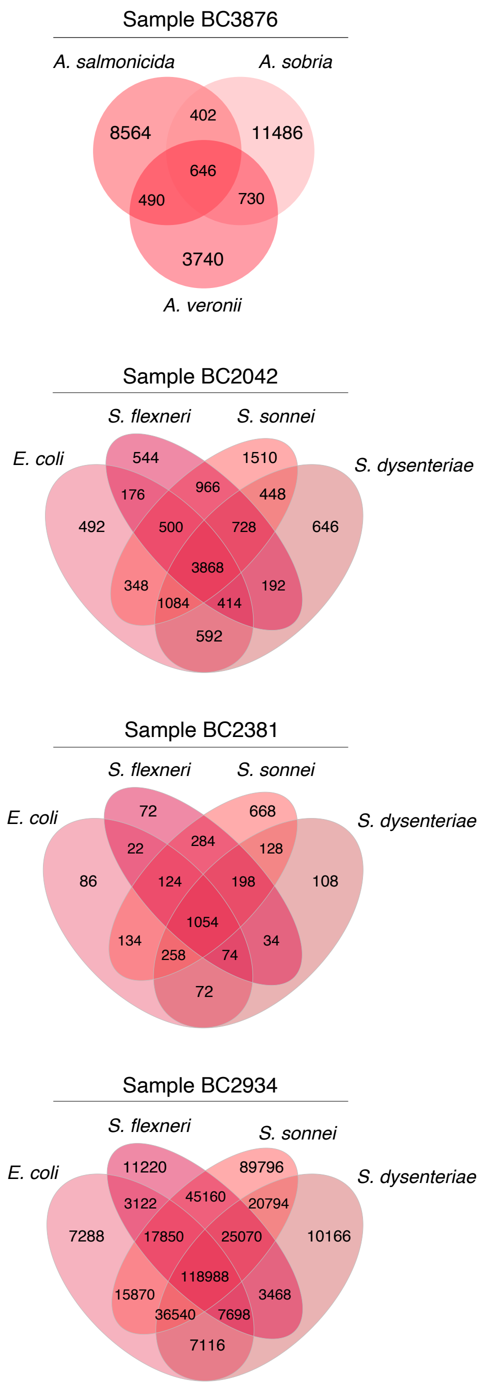


**Figure S4.** Venn diagram comparing reads assigned to species of *Aeromonas* and the *Escherichia coli-Shigella* group in a single sample, showing shared reads (intersections) and unique reads (non-intersecting areas) for each species.
